# Element accumulation in the tracheal and bronchial cartilages of monkeys

**DOI:** 10.1101/2024.03.01.582912

**Authors:** Cho Azuma, Takao Oishi, Yoshiyuki Tohno, Lining Ke, Xiao-zhen Zhao, Takeshi Minami, Noriko Horii-Hayashi, Koichi Inoue

**Affiliations:** Department of Anatomy and Cell Biology, Nara Medical University, Kashihara, Nara, Japan; Systems Neuroscience Section, Center for the Evolutionary Origins of Human Behaviors, Kyoto University, Inuyama, Aichi, Japan; Department of Human Anatomy and Histo-Embryology, School of Basic Medical Sciences, Fujian Medical University, Fuzhou, China; Laboratory of Environmental Biology, Department of Life Science, Faculty of Science and Technology, Kinki University, Higashi-Osaka, Osaka, Japan

## Abstract

Compositional changes in the tracheal and bronchial cartilages can affect respiratory ventilation and lung function. We aimed to elucidate element accumulation in the tracheal and bronchial cartilages of monkeys and divided it into four sites: tracheal, tracheal bifurcation, left bronchial, and right bronchial cartilages. The elemental content was analyzed using inductively coupled plasma atomic emission spectrometry.

The average calcium content was two to three times higher in the tracheal cartilage than in the other three cartilages. The trends of phosphorus and zinc were similar to those of calcium. The average calcium, phosphorus, and zinc contents were the highest in the tracheal cartilage and decreased in the following order: the left bronchial, right bronchial, and tracheal bifurcation cartilages. These findings revealed that differences existed in element accumulation between different sites within the same airway cartilage and that calcium, phosphorus, and zinc accumulation mainly occurred in the tracheal cartilage.

A substantial direct correlation was observed between age and calcium content in the tracheal and bronchial cartilages and all such monkeys with high calcium content were > four years of age. These results suggest that calcium accumulation occurs in the tracheal and bronchial cartilages after reaching a certain age.

An extremely substantial direct correlation was observed between calcium and phosphorus contents in the tracheal and bronchial cartilages. This finding is similar to the previously published calcium and phosphorus correlations in several other cartilages, suggesting that the calcium and phosphorus contents of cartilage exist in a certain ratio.

## Introduction

The lumen must always remain open since the trachea and bronchi are airways through which air continuously enters and leaves. The trachea and bronchi have C-shaped hyaline cartilages on the anterior and lateral walls. Furthermore, the cartilage rings have sufficient strength to play a crucial role in maintaining the open state of the respiratory tract to prevent trachea and bronchi collapse and obstruction during respiration. Compositional changes in the tracheal and bronchial cartilages can affect respiratory ventilation and lung function. For example, calcification and ossification can make surgical airway provision, which involves tracheal cartilage incision and intubation, challenging [1-2]. Since these cases are likely to increase with the advancing aging population, studying compositional changes in the tracheal and bronchial cartilages is extremely useful.

Several reports on the calcification, ossification, and stiffness of the tracheal and bronchial cartilages exist using X-ray, computed tomography (CT), von Kossa staining, and mechanical tests. Kusafuka et al. [3] studied tracheal cartilage ossification in aged humans using histological and immunohistochemical analyses. Ohkubo et al. [4-5] investigated the CT findings of benign tracheobronchial lesions with calcification. Liu et al. [6] reported that calcification was biochemically and histologically observed in the tracheal cartilage of rabbits. Sasano et al. [7] studied the calcification process during rat tracheal cartilage development. Yokoyama et al. [8] reported a case of tracheobronchial stenosis with calcification. Furthermore, Safshekan et al. [9] reported that tracheal cartilage stiffness increased with age. However, the compositional changes using direct chemical analysis and comparing the calcification incidence in the four regions of the monkey tracheal and bronchial cartilages have not yet been investigated. Furthermore, no clinical studies have compared calcification incidence at these four sites. Considering the morphological and genetic similarities between monkeys and humans, the trend of metal retention in monkeys if proven similar to that in humans could be used for basic research on tracheal cartilage calcification and pre-clinical studies, such as in the development of new instrumentation [10-11]. Therefore, the authors investigated compositional changes, different element accumulations, and age-related changes in various parts of the tracheal and bronchial cartilages.

## Materials and methods

### Sampling

All animal experiments were performed in accordance with the US National Institutes of Health Guide for the Care and Use of Laboratory Animals and the Guidelines for Care and Use of Nonhuman Primates (Ver. 3, 2010; Primate Research Institute, Kyoto University, Japan). The study protocol was approved by the Animal Welfare and Animal Care Committee of the Primate Research Institute, Kyoto University (Permission No. 2010-071, 2011-019, 2012-029, 2013-024, 2014-028, 2015-120, 2016-045, 2018-030). The monkeys were bred at the Primate Research Institute of Kyoto University. They were pretreated with an intramuscular injection of ketamine hydrochloride (10 mg/kg) and deeply anesthetized via intravenous pentobarbital sodium administration (Nembutal, 30 mg/kg). The monkeys were subsequently perfused through the left ventricle with ice-cold saline (0.5 L of ice-cold saline containing 2 mL (2,300 U) of heparin sodium, followed by 1–2 L of ice-cold fixative consisting of 2% paraformaldehyde and 0.5% glutaraldehyde in 0.15 M phosphate buffer (pH 7.4). After perfusion, the tracheal and bronchial cartilages were resected from the monkeys. The tracheal and bronchial cartilages were further separated into four sites: tracheal cartilage (TC), tracheal bifurcation cartilage (TBC), left bronchial cartilage (LBC), and right bronchial cartilage (RBC) (Fig 1) to investigate the differences between different sites. The rhesus and Japanese monkeys consisted of 13 males and 4 females, ranging in age from 0.1 to 29 years.

**Fig 1.**
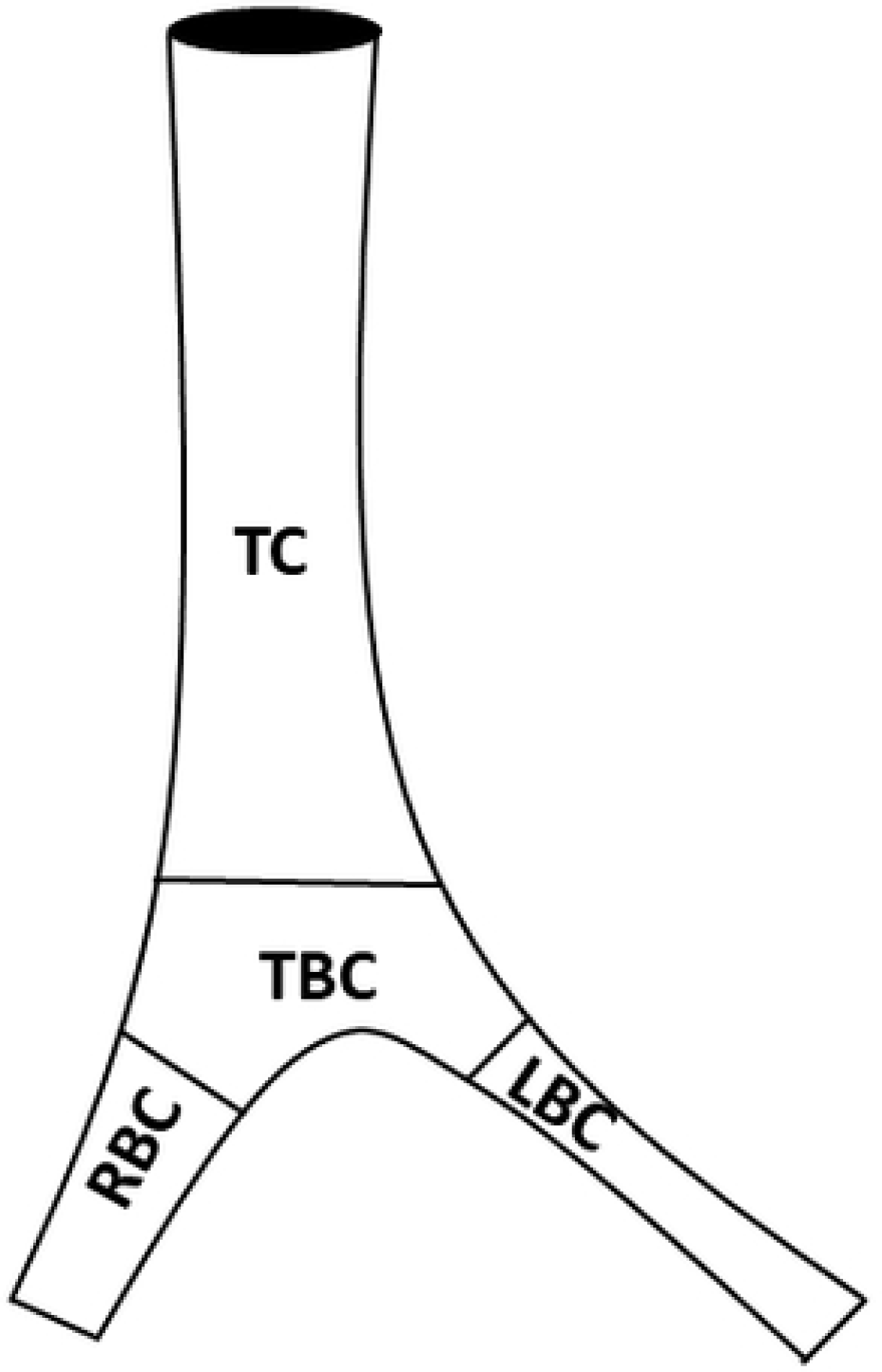
Sites of tracheal cartilage (TC), tracheal bifurcation cartilage (TBC), left bronchial cartilage (LBC), and right bronchial cartilage (RBC).

### Determining the elements

The monkey tracheal and bronchial cartilages were thoroughly washed with distilled water and dried at 80°C for 16 h. After adding 1 mL of concentrated nitric acid to the samples to incinerate, they were heated at 100°C for 2 h in a dry-block bath (FS-620; Tokyo, Japan). After adding concentrated perchloric acid (0.5 mL), the samples were heated at 100°C for an additional 2 h. The samples were subsequently adjusted to a 10 mL volume by adding ultrapure water and filtering through a filter paper (No. 7, Toyo Roshi, Osaka, Japan). The resulting filtrates were analyzed using an inductively coupled plasma atomic emission spectrometer (ICPS-7510, Shimadzu, Kyoto, Japan). The conditions were 1.2 kW of power from a radiofrequency generator, a plasma argon flow rate of 1.2 L/min, a cooling gas flow of 14 L/min, a carrier gas flow of 1.0 L/min, an entrance slit of 20 µm, an exit slit of 30 µm, an observation height of 15 mm, and an integration time lapse of 5 s. The amounts of elements were expressed on a dry-weight basis.

### Statistical analyses

Statistical analyses were performed using GraphPad Prism (version 3.0; GraphPad Software, San Diego, CA, USA). The association between the parameters was investigated using Pearson’s correlation coefficient. One-way analysis of variance (ANOVA) followed by the Student–Newman–Keuls test was used to compare the differences among the four groups. Statistical significance was set at *p*-value < 0.05. Data are expressed as the mean ± standard deviation.

## Results

### Elemental contents

Table 1 lists the average contents of the elements in the TC, TBC, LBC, and RBC.

**Table 1.** Average contents of elements in the TC, TBC, LBC, and RBC.

|  | Ca (mg/g) | P (mg/g) | Mg (mg/g) | S (mg/g) | Zn (μg/g) | Fe (μg/g) | Na (μg/g) |
| --- | --- | --- | --- | --- | --- | --- | --- |
| **TC** | 39.98 ± 35.78 | 17.03 ± 17.03 | 1.555 ± 1.451 | 7.809 ± 1.966 | 524.5 ± 360.8 | 277.1 ± 179.7 | 522.8 ± 346.7 |
| **TBC** | 13.07 ± 14.51 | 4.79 ± 6.50 | 1.306 ± 1.018 | 6.666 ± 0.962 | 216.1 ± 136.7 | 244.3 ± 213.3 | 402.3 ± 416.0 |
| **LBC** | 18.87 ± 21.81 | 7.51 ± 9.75 | 1.395 ± 1.094 | 6.816 ± 1.370 | 302.1 ± 182.4 | 321.8 ± 337.4 | 410.2 ± 486.6 |
| **RBC** | 13.97 ± 19.56 | 5.46 ± 9.03 | 1.362 ± 1.032 | 6.673 ± 1.206 | 233.2 ± 216.1 | 263.3 ± 259.6 | 412.6 ± 524.3 |

The average calcium content was two to three times higher in the TC than in the TBC, LBC, and RBC. One-way ANOVA revealed statistically significant differences among the TC, TBC, LBC, and RBC (*p* = 0.0057). Further analysis using the Student–Newman– Keuls test revealed that the TC was significantly higher than the TBC (*p* = 0.0101), LBC *(p* = 0.0135), and RBC (*p* = 0.0074). However, no significant differences were observed among the TBC, LBC, and RBC.

The average phosphorus content was two to three times higher in the TC than in the TBC, LBC, and RBC. One-way ANOVA revealed statistically significant differences among the TC, TBC, LBC, and RBC (*p* = 0.0079). Further analysis using the Student– Newman–Keuls test indicated that the TC was significantly higher than the TBC (*p* = 0.0124), LBC *(p* = 0.0166), and RBC (*p* = 0.0109). However, no significant differences were observed among the TBC, LBC, and RBC.

The average zinc content was approximately two times higher in the TC than in the TBC, LBC, or RBC. One-way ANOVA revealed statistically significant differences among the TC, TBC, LBC, and RBC (*p* = 0.0012). Further analysis using the Student– Newman–Keuls test revealed that the TC was significantly higher than the TBC (*p* = 0.0021), LBC (*p* = 0.0086), and RBC (*p* = 0.0021). However, no significant differences were observed among the TBC, LBC, and RBC.

The average sulfur, magnesium, iron, and sodium contents in the TC were almost the same as those in the TBC, LBC, and RBC. No significant differences were observed among the TC, TBC, LBC, and RBC.

### Age-related changes in the elements

Figure 2 illustrates the age-related changes in the calcium content of TC, TBC, LBC, and RBC. The correlation coefficients between age and calcium content were estimated to be 0.497 (*p* = 0.043), 0.885 (*p* < 0.0001), 0.825 (*p* < 0.0001), and 0.882 (*p* < 0.0001) for TC, TBC, LBC, and RBC, respectively. A significant direct correlation was observed between age and calcium content in the TC, and extremely significant direct correlations were observed between age and calcium content in the TBC, LBC, and RBC. The calcium content suddenly increased at approximately four years of age in the TC, whereas it progressively increased in the other three cartilages with age. Notably, all monkeys with high calcium content in the TC, TBC, LBC, and RBC were > 4 years of age.

**Fig 2.**
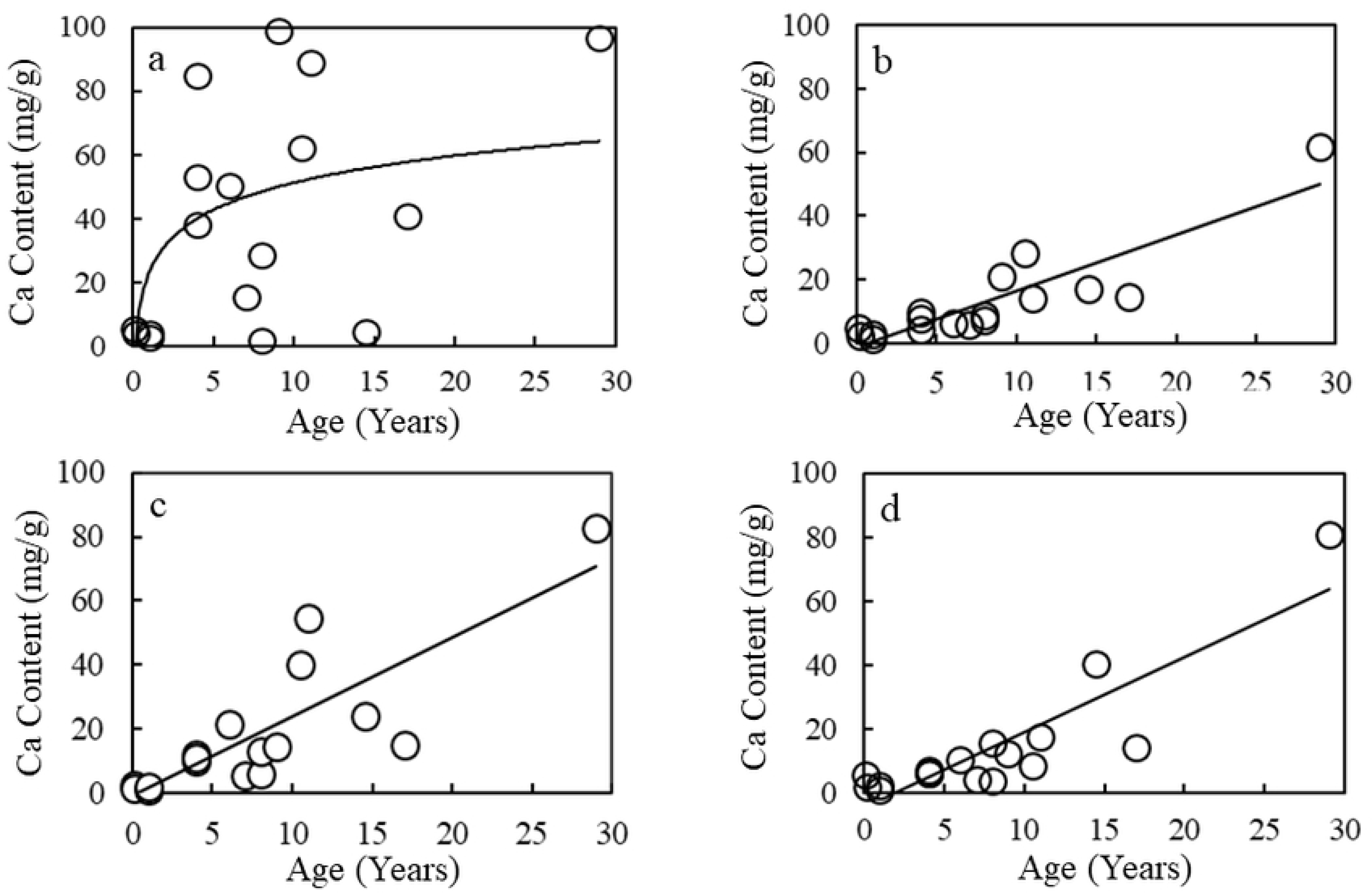
Age-related changes in the calcium contents in the TC (a), TBC (b), LBC (c), and RBC (d).

Figure 3 illustrates the age-related changes in the phosphorus content of TC, TBC, LBC, and RBC. The correlation coefficients between age and phosphorus content were estimated to be 0.450 (*p* = 0.070), 0.832 (*p* < 0.0001), 0.795 (*p* = 0.0001), and 0.844 (*p* < 0.0001) for TC, TBC, LBC, and RBC, respectively. Extremely significant direct correlations were observed between age and phosphorus content in the TBC, LBC, and RBC but not in the TC. The phosphorus content suddenly increased at approximately four years of age in the TC, whereas it progressively increased in the other three cartilages with age. Notably, all monkeys with high phosphorus content in the TC, TBC, LBC, and RBC were > 4 years of age.

**Fig 3.**
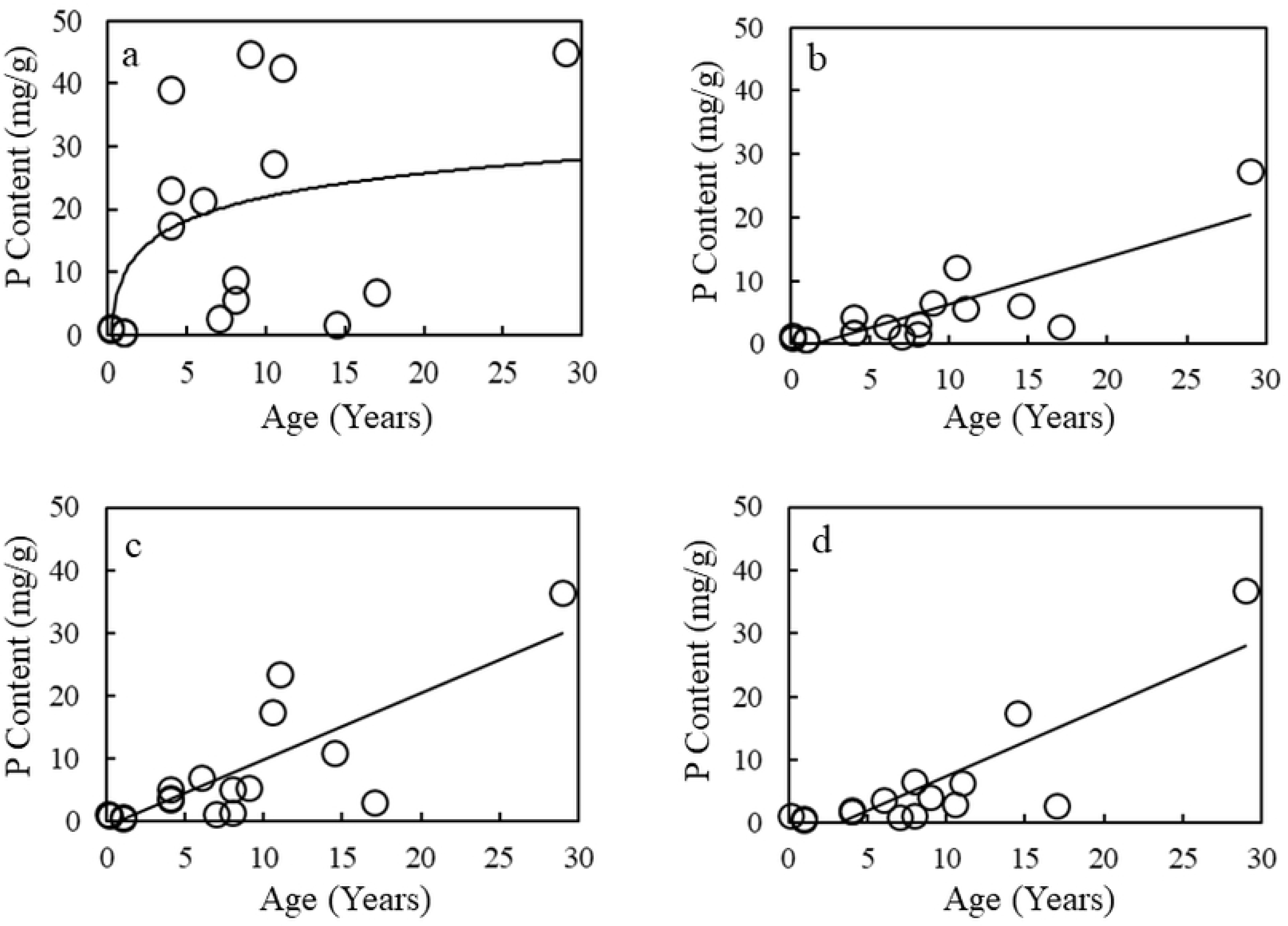
Age-related changes in the phosphorus content in the TC (a), TBC (b), LBC (c), and RBC (d).

Figure 4 illustrates the age-related changes in the magnesium contents in the TC, TBC, LBC, and RBC. The correlation coefficients between age and magnesium content were estimated to be 0.317 (*p* = 0.216), 0.436 (*p* = 0.080), 0.515 (*p* = 0.034), and 0.513 (*p* = 0.035) for TC, TBC, LBC, and RBC, respectively. Significant direct correlations were observed between age and magnesium content in the LBC and RBC; however, the correlations in the TC and TBC were statistically insignificant.

**Fig 4.**
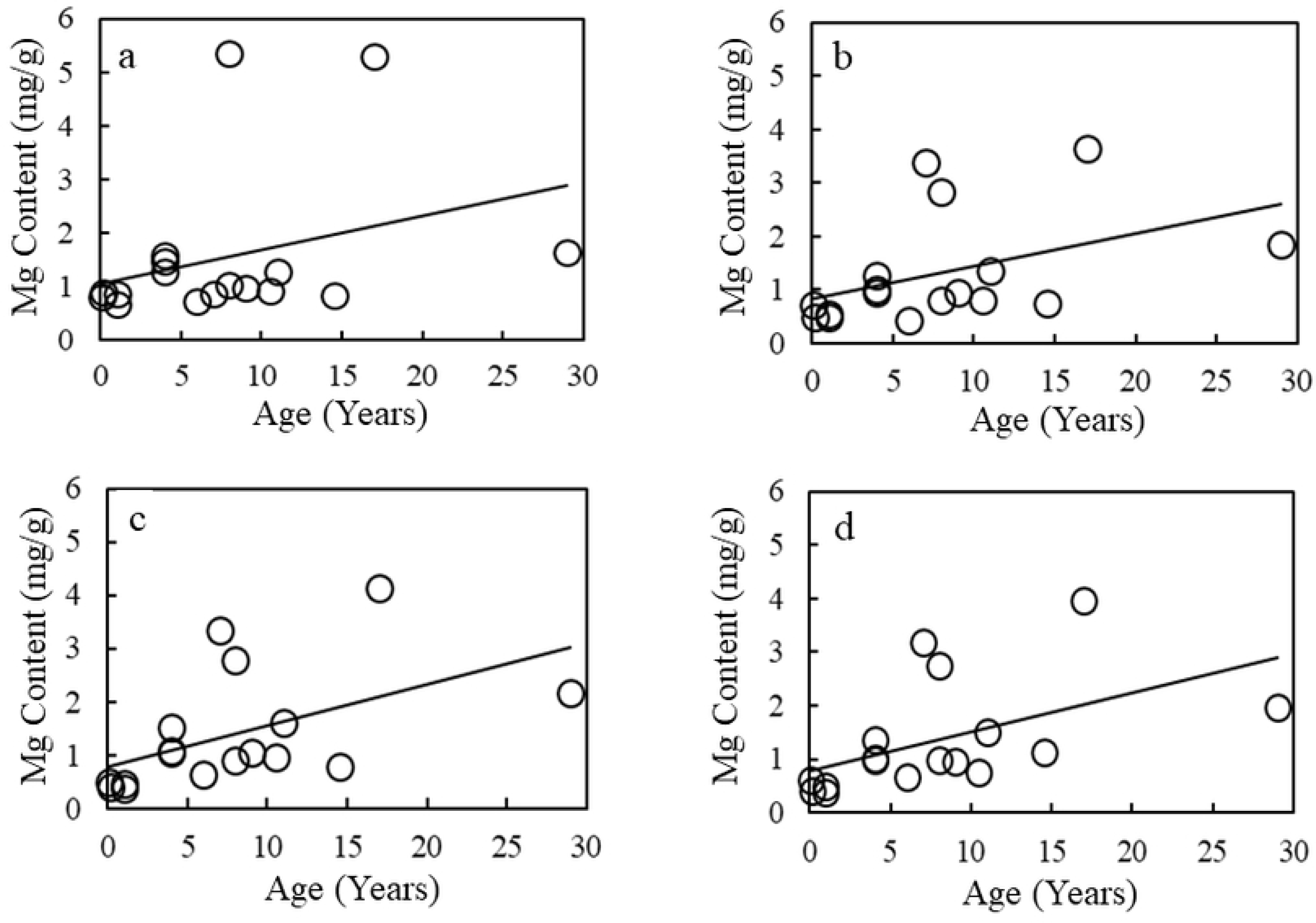
Age-related changes in the magnesium content in the TC (a), TBC (b), LBC (c), and RBC (d).

Figure 5 illustrates the age-related changes in the zinc contents of TC, TBC, LBC, and RBC. The correlation coefficients between age and zinc content were estimated to be 0.344 (*p* = 0.177), 0.483 (*p* = 0.050), 0.481 (*p* = 0.051), and 0.608 (*p* = 0.010) for TC, TBC, LBC, and RBC, respectively. A significant direct correlation was observed between age and zinc content in the RBC; however, no significant correlations were observed with TC, TBC, or LBC.

**Fig 5.**
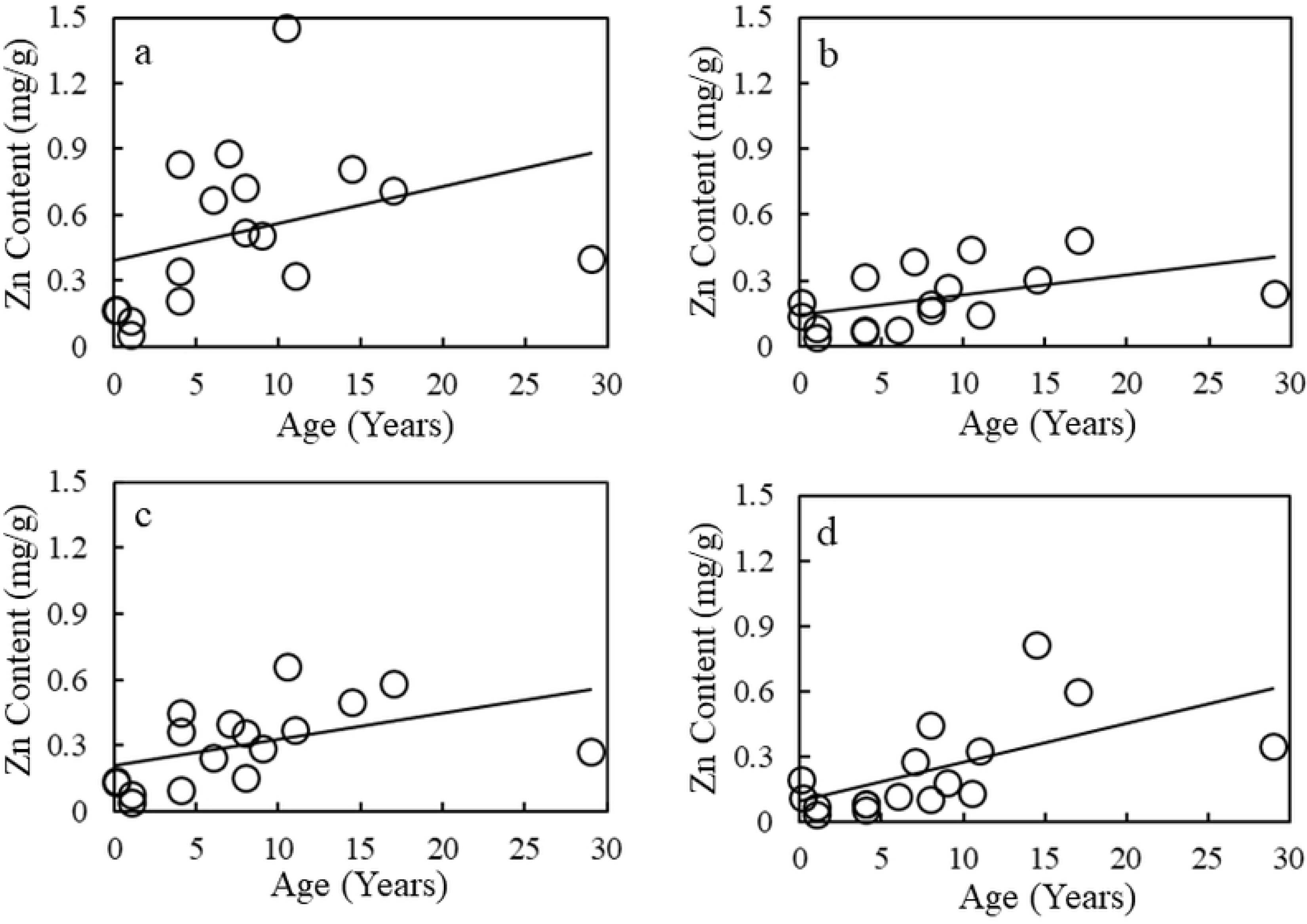
Age-related changes in the zinc content in the TC (a), TBC (b), LBC (c), and RBC (d).

### Relationships of the elemental contents between the TC and the other three cartilages

Figure 6 illustrates the relationship of the calcium content between the TC and the other three cartilages. The correlation coefficients were estimated to be 0.611 (*p* = 0.009) between TC and TBC, 0.669 (*p* = 0.003) between TC and LBC, and 0.410 (*p* = 0.102) between TC and RBC. Extremely significant direct correlations in the calcium content were observed between TC and either TBC or LBC but not between TC and RBC.

**Fig 6.**
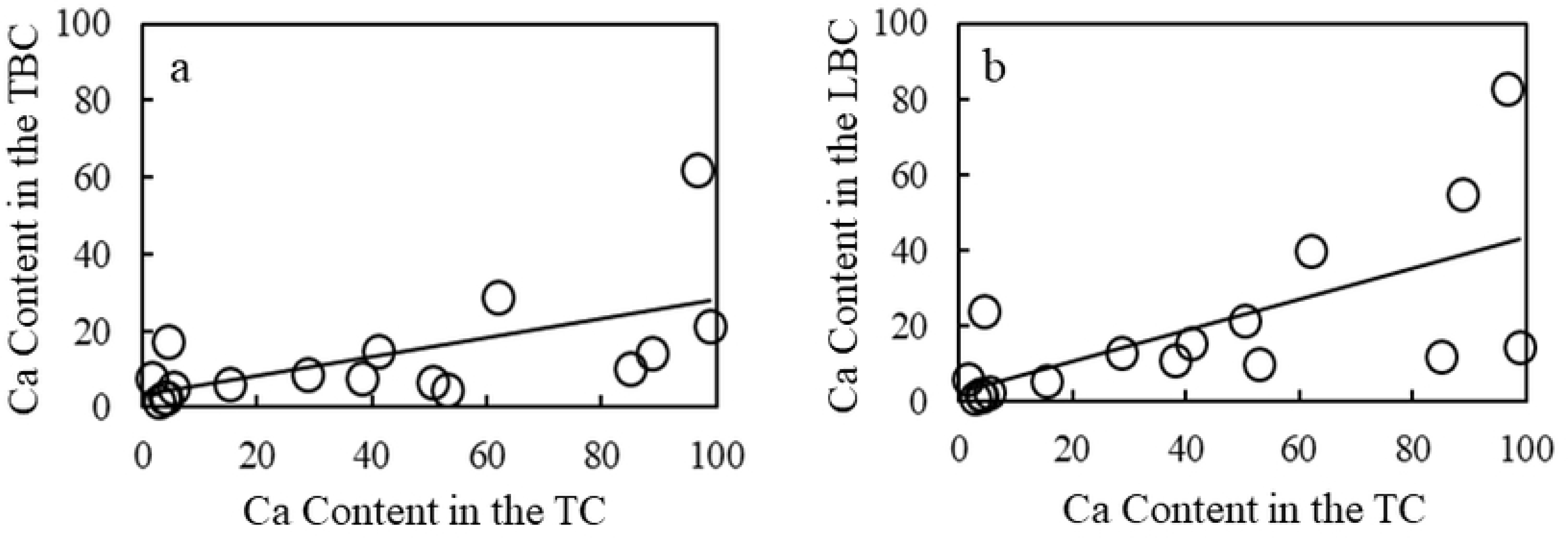
Relationships of the calcium content between the TC and either TBC (a) or LBC (b).

Figure 7 illustrates the relationship of the phosphorus content between the TC and the other three cartilages. The correlation coefficients were estimated to be 0.614 (*p* = 0.009) between TC and TBC, 0.671 (*p* = 0.003) between TC and LBC, and 0.394 (*p* = 0.118) between TC and RBC. Extremely significant direct correlations in the phosphorus content were observed between TC and either TBC or LBC but not between TC and RBC.

**Fig 7.**
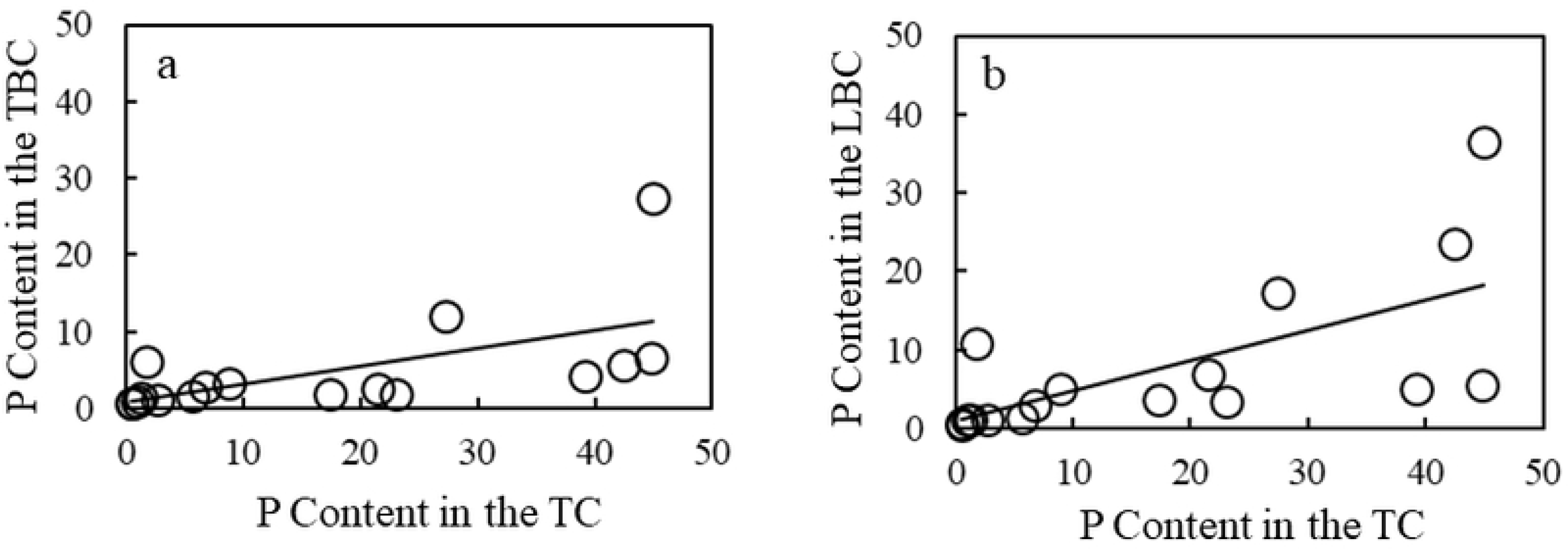
Relationships of the phosphorus content between the TC and either TBC (a) or LBC (b).

Figure 8 illustrates the relationship of the magnesium content between the TC and the other three cartilages. The correlation coefficients were estimated to be 0.731 (*p* = 0.0009) between TC and TBC, 0.742 (*p* = 0.0006) between TC and LBC, and 0.751 (*p* = 0.0005) between TC and RBC. Extremely significant direct correlations in the magnesium content were observed between the TC and the other three cartilages.

**Fig 8.**
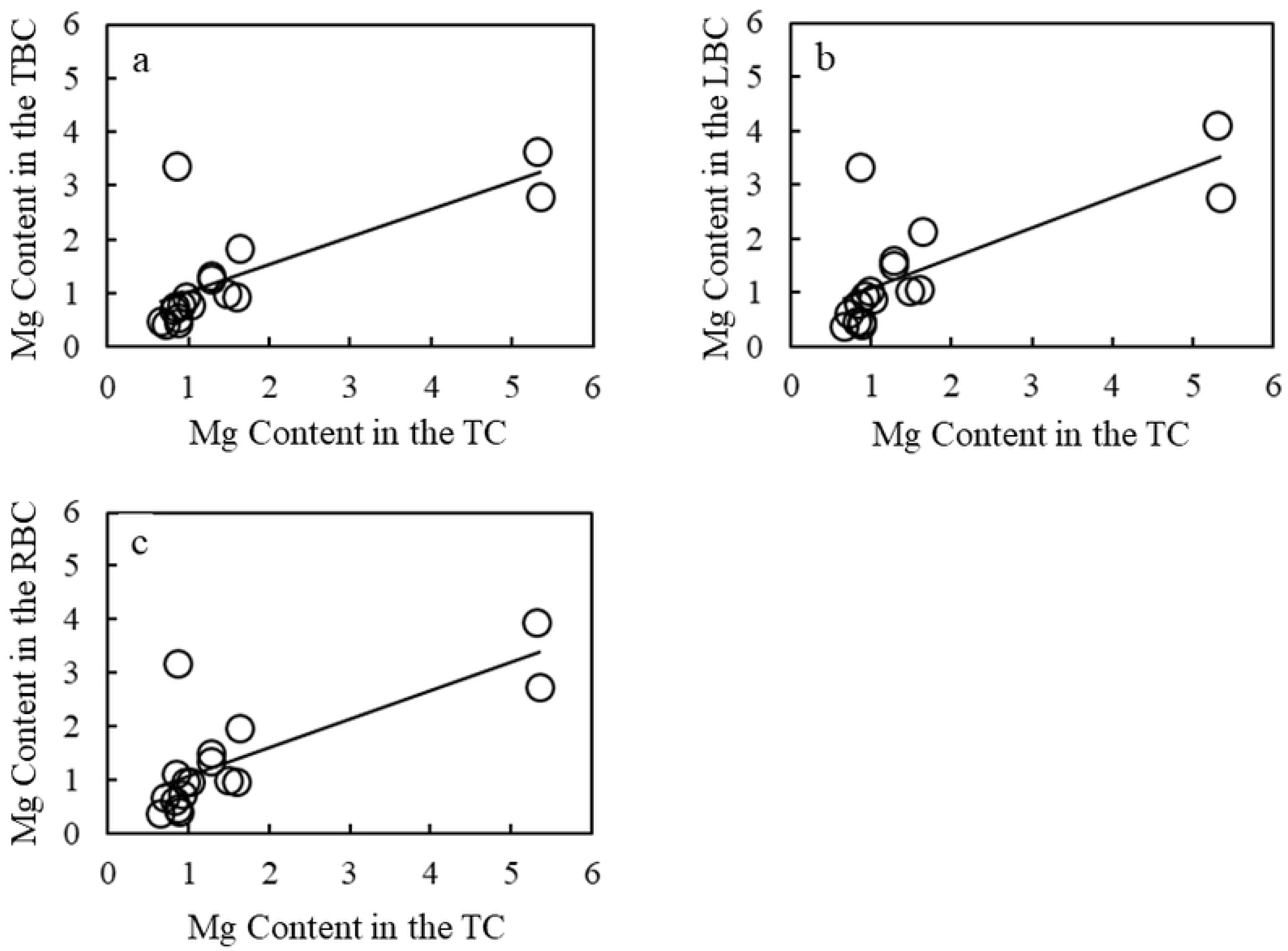
Relationships of the magnesium content between the TC and TBC (a), LBC (b), and RBC (c).

Figure 9 illustrates the relationship of the zinc content between the TC and the other three cartilages. The correlation coefficients were estimated to be 0.618 (*p* = 0.008) between TC and TBC, 0.815 (*p* < 0.0001) between TC and LBC, and 0.322 (*p* = 0.207) between TC and RBC. Extremely significant direct correlations in the zinc content were observed between TC and either TBC or LBC but not between TC and RBC.

**Fig 9.**
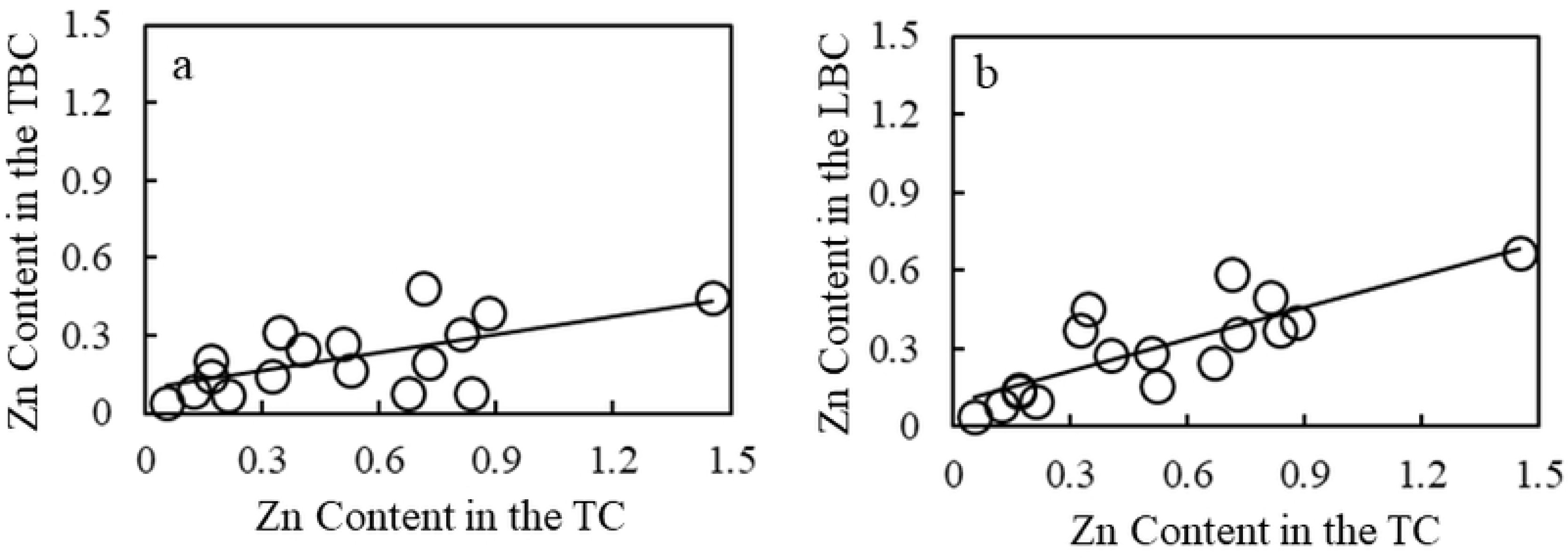
Relationships of the zinc content between the TC and either TBC (a) or LBC (b).

Regarding the sulfur, iron, and sodium contents, no significant correlations were observed between the TC and the other three cartilages.

### Relationships among elements

Table 2 lists the relationships among the seven elements in the TC. An extremely significant direct correlation was observed between the calcium and phosphorus contents in the TC. The correlations between the calcium and magnesium contents and between the phosphorus and magnesium contents in the TC were statistically insignificant. Regarding the TBC, LBC, and RBC, similar results were obtained.

**Table 2.** Relationships among seven elements in the TC.

|  | Correlation Coefficient and *p* Value |  |  |  |  |  |
| --- | --- | --- | --- | --- | --- | --- |
| Element | P | Mg | S | Zn | Fe | Na |
| Ca | 0.979 (<0.0001) | -0.065 (0.803) | 0.226 (0.382) | 0.178 (0.494) | -0.110 (0.675) | 0.256 (0.321) |
| P |  | -0.103 (0.694) | 0.260 (0.314) | 0.119 (0.650) | -0.142 (0.588) | 0.242 (0.349) |
| Mg |  |  | 0.146 (0.576) | 0.095 (0.717) | -0.006 (0.981) | -0.056 (0.832) |
| S |  |  |  | -0.172 (0.510) | 0.027 (0.918) | 0.311 (0.225) |
| Zn |  |  |  |  | 0.617 (0.008) | -0.216 (0.405) |
| Fe |  |  |  |  |  | -0.442 (0.076) |

## Discussion

While tracheal cartilage ossification and calcification develop in various diseases, they also occur with age [2]. They often cause stiffness and/or stenosis and clinical problems during medical procedures such as intubation and tracheostomy. Although their extent can be detected using CT and other devices, measuring the actual amount of each element and corroborating this information as evidence is still meaningful [12-14].

This study revealed that the average calcium content was two to three times higher in the TC than in the TBC, LBC, and RBC. Considerable differences in the average calcium content were observed between the TC and the other three cartilages. The changing trends in the average phosphorus and zinc contents were parallel to that of the average calcium content. These results are reasonable because phosphorus is present as phosphate in ossified tissues following calcium accumulation and contributes to the structure and function, and zinc plays important roles in collagen synthesis and the activity of alkaline phosphatases that produce hydroxyapatite [15, 16].

Cartilage is divided into three classes: hyaline, fibro, and elastic. The tracheal and bronchial cartilages are classified as hyaline. The authors previously investigated compositional changes with age in human hyaline cartilages, such as the trachea [17], xiphoid process [18], and costal cartilage [18]; human fibrocartilages, such as the articular disk of the temporomandibular joint [19], meniscus [20], pubic symphysis [21], and intervertebral disk [22]; and human elastic cartilages, such as the epiglottal cartilage [23]. A high calcium accumulation sometimes occurred in the trachea, xiphoid process, costal cartilage, pubic symphysis, and intervertebral disk but not in the meniscus, articular disk of the temporomandibular joint, and epiglottal cartilage. This study found that a high calcium accumulation sometimes occurred in the TC, TBC, LBC, and RBC of monkeys. Some reports regarding the calcification or ossification of tracheal and bronchial cartilages exist in humans [3-5, 24-25], rabbits [6], and rats [7] that used X-ray, CT, and von Kossa staining. Furthermore, the authors previously investigated compositional changes in the human trachea and found that a high calcium accumulation often occurs in the human trachea [17]. These results are consistent with our results, which demonstrate that high calcium accumulation occurs in monkey tracheal and bronchial cartilages.

Within the human coronary artery [26], plantar aponeurosis [27], and palmar aponeurosis [27], significant differences have been found in elemental accumulation between different sites, even within the same organ. The current study found that the average calcium content of the cartilage was the highest in the TC and decreased in the following order: LBC, RBC, and TBC. Substantial differences in the average calcium content were observed between the TC and the other three cartilages. The changing trends in the average phosphorus and zinc contents paralleled with those of the average calcium content. The average phosphorus and zinc contents of the cartilage were the highest in TC and decreased in the following order: LBC, RBC, and TBC. Considerable differences in the average phosphorus and zinc contents were observed between the TC and the other three cartilages. Therefore, differences are likely to exist in the elemental accumulation between different sites within the same airway cartilage.

Generally, the age of rhesus and Japanese monkeys multiplied by three is believed to correspond to human age. Ohkubo et al. [5] reported that calcification started to appear in the second decade of life in the human tracheal cartilage. Liu et al. (6) reported that calcification started after 15 weeks in the tracheal cartilage of rabbits. Sasano et al. [7] reported that calcification occurred in the tracheal cartilage of rats 10 weeks after birth. Notably, all monkeys in our study with high calcium content in the tracheal and bronchial cartilages were > four years of age (corresponding to 12 years in humans). Furthermore, a substantial direct correlation was observed between age and calcium content in the tracheal and bronchial cartilage. Thus, these results suggest that calcium accumulation mainly occurs in the tracheal and bronchial cartilage of humans and animals after reaching a particular age.

Elucidating age-related compositional changes in tissues and organs is challenging in humans. Monkeys were selected as the research subjects because approximately 21 bronchial cartilage rings of hyaline cartilage in monkeys [28] are similar to the 18–22 bronchial cartilage rings of the hyaline cartilage in humans [29], and monkey specimens of various ages can be collected. Rhesus monkeys and Japanese monkeys have almost the same tracheal length and the number of C-shaped tracheal cartilage rings [28]. Therefore, the tracheal cartilages of the rhesus and Japanese monkeys were studied in the same manner. The authors previously investigated age-related changes in elements by direct chemical analysis of the monkey cardiac walls [30], sinoatrial node [31], cardiac valves [32], tendon of the peroneus longus muscle [33], ligamentum capitis femoris [34], and various arteries [35-40] and found that the elements did not uniformly accumulate in various monkey tissues and organs with age. In the cardiac walls, sinoatrial nodes, cardiac valves, and coronary artery [35-36], the calcium content gradually decreased with development. Conversely, in most arteries [37-40] and the tendon of the peroneus longus muscle, the calcium content progressively increased with age. These results suggest that calcium accumulation in monkey tissues and organs has two completely opposite trends of increasing or decreasing with age. In other words, changes in the calcium levels in monkey tissues and organs with age were divided into calcium-accumulated and calcium-released types. The present study revealed that the average calcium content progressively increased with age in the monkey tracheal and bronchial cartilages. Therefore, changes in the calcium levels in the monkey tracheal and bronchial cartilages were classified as the calcium-accumulated type with age.

Considerable direct correlations in calcium, phosphorus, magnesium, and zinc contents were observed between TC and either TBC or LBC. Therefore, calcium, phosphorus, magnesium, and zinc compositions in TC, TBC, and LBC are possibly closely related. However, the correlation between the calcium, phosphorus, and zinc contents of TC and RBC was statistically insignificant, which warrants further investigation.

The authors [18-23] investigated the elemental contents in seven cartilage types: the articular disk of the temporomandibular joint, costal cartilage, epiglottal cartilage, intervertebral disk, left medial meniscus, pubic symphysis, and xiphoid process. The following findings were obtained: a considerable direct correlation between calcium and phosphorus contents in these cartilages was observed, except for the articular disk of the temporomandibular joint. However, no substantial direct correlations between the calcium and magnesium contents in the three cartilage types or between phosphorus and magnesium contents in the five cartilage types were found. Furthermore, a significant direct correlation existed between calcium and phosphorus contents in the TC, TBC, LBC, and RBC; however, no significant correlations between calcium and magnesium contents or between phosphorus and magnesium contents were observed. These findings suggest that the calcium and phosphorus contents of cartilage exist in a certain ratio. In the arteries, significant direct correlations existed between calcium, phosphorus, and magnesium contents [40]. The differences in the relationships between calcium, phosphorus, and magnesium contents in the arteries and cartilages warrant further investigation.

## Conclusions

The average calcium content was the highest in the tracheal cartilage and decreased in the following order: left bronchial, right bronchial, and tracheal bifurcation cartilages. Changes in calcium accumulation in the tracheal and bronchial cartilages were age-related and occurred after reaching a certain age. These results provide meaningful basic evidence supporting age-related clinical airway problems.

## Acknowledgments

This study was supported by the Cooperation Research Program (2010-2016, 2018) of the Primate Research Institute of Kyoto University.

